# Rational Design of a Flavoenzyme for Aerobic Nicotine Catabolism

**DOI:** 10.1101/2024.07.11.603087

**Authors:** Haiyang Hu, Zhaoyong Xu, Zhiyao Zhang, Peizhi Song, Frederick Stull, Ping Xu, Hongzhi Tang

## Abstract

Enzymatic therapy with nicotine-degrading enzyme is a new strategy in treating nicotine addiction, which can reduce nicotine concentrations and weaken withdrawal in the rat model. However, when O_2_ is used as the electron acceptor, no satisfactory performance has been achieved with one of the most commonly studied and efficient nicotine-catabolizing enzymes, NicA2. To obtain more efficient nicotine-degrading enzyme, we rationally designed and engineered a flavoenzyme Pnao, which shares high structural similarity with NicA2 (RMSD = 1.143 Å) and efficiently catalyze pseudooxynicotine into 3-succinoyl-semialdehyde pyridine using O_2_. Through amino acid alterations with NicA2, five Pnao mutants were generated, which can degrade nicotine in Tris-HCl buffer and retained catabolic activity on its natural substrate. Nicotine-1’-*N*-oxide was identified as one of the reaction products. Four of the derivative mutants showed activity in rat serum and Trp220 and Asn224 were found critical for enzyme specificity. Our findings offer a novel avenue for research into aerobic nicotine catabolism and provides a promising method of generating additional nicotine-catalytic enzymes.

## Introduction

Tobacco use seriously harms nearly every organ in the human body, causing cardiovascular disease, chronic obstructive pulmonary disease, lung cancer, and other complicated disorders. It kills approximately seven million people each year (1), far exceeding deaths due to alcohol and illicit drug use (2). During pregnancy, the risk of stillbirth, neonatal death, and perinatal death increases along with the amount of tobacco smoked (3). Furthermore, cognitive differences and behavioral-control disorders are seen in primary school children who have prenatal or postnatal smoke exposure (4). Therapy for nicotine addiction facilitates in smoking cessation because nicotine is the major active ingredient in tobacco that results in addiction and various diseases such as atherosclerosis, lung cancer, and cardiac dysfunction (5–8). In recent years, an approach to smoking cessation has emerged that involves decreasing the nicotine concentration in the peripheral circulation through catabolism of nicotine to prevent it from accessing the brain. This novel enzymatic approach is primarily based on the nicotine oxidoreductase NicA2 (9–11).

NicA2 is a vital enzyme in *Pseudomonas putida* S16, a strain that can effectively mineralize nicotine via the pyrrolidine pathway (12). In this pathway, the enzymes NicA2, Pnao, and Sapd are respectively responsible for following conversions: nicotine to *N*-methylmyosmine (NMM), pseudooxynicotine (PN) to 3-succinoyl-semialdehyde pyridine (SAP), and SAP to 3-succinoyl pyridine (SP); NMM can be spontaneously hydrolyzed to PN (Fig. 1a). SP is finally transformed to fumaric acid which is an intermediate metabolite in the tricarboxylic acid (TCA) cycle (13).

**Fig. 1.**
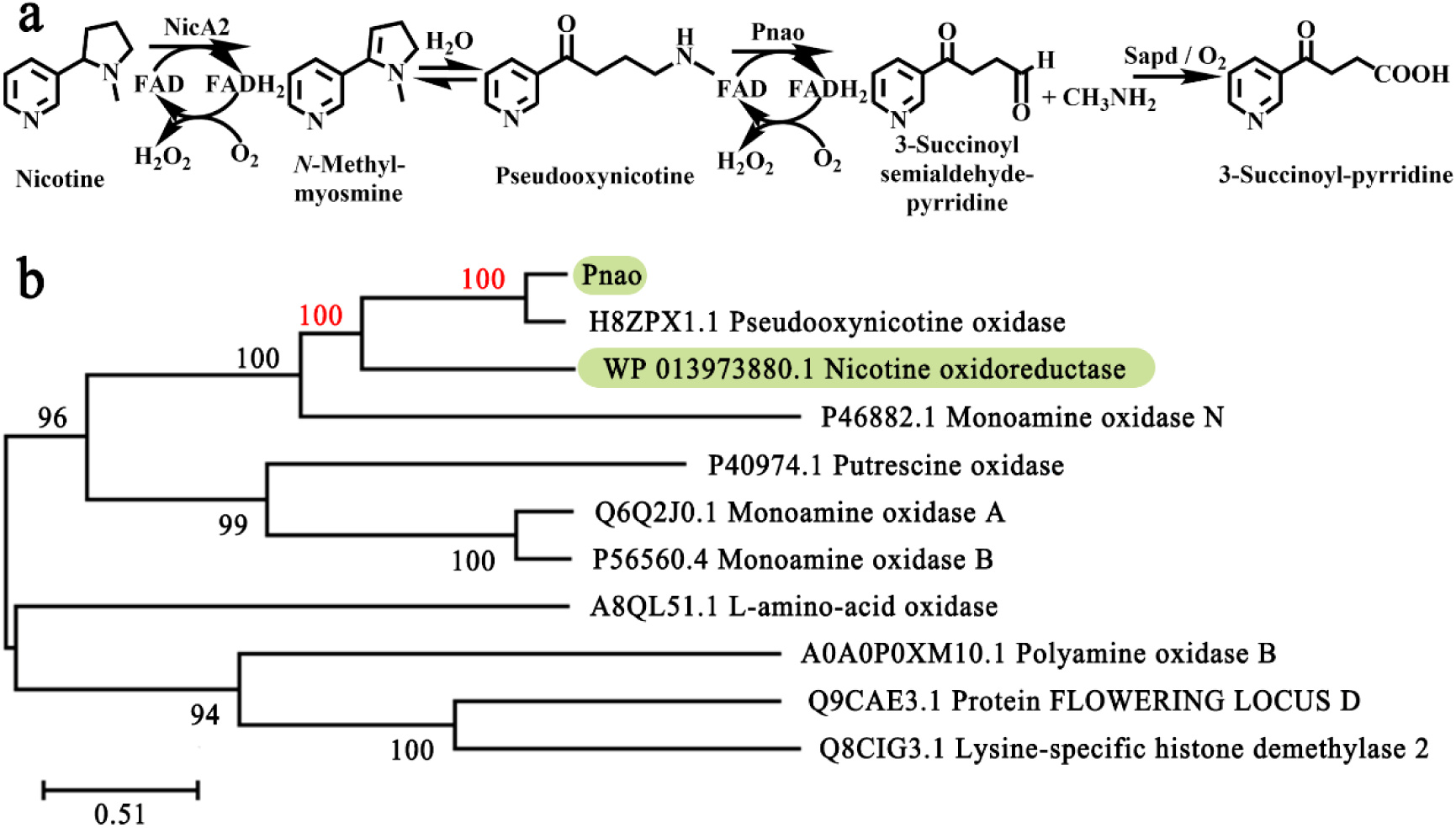
Pnao sequence and biochemical pathway involvement. (**a**) NicA2 converts nicotine to *N*-methylmyosmine (NMM), which is spontaneously hydrolyzed to pseudooxynicotine (PN) before being oxidized to 3-succinoyl-semialdehyde pyridine (SAP) by Pnao. O_2_ acts as the electron acceptor in the absence of cytochrome *c*. SAP can be further oxidized to 3-succinoyl pyridine (SP) by Sapd or O_2_. **(b)** Phylogenetic tree showing Pnao and related enzymes.

NicA2 has long been targeted in attempts to increase the efficiency of nicotine enzymatic catabolism. Although it is the most well studied and efficient nicotine degrader, the *k*_cat_ of NicA2 for nicotine is only 6.2×10^−3^ s^−1^ (14). Several mutations have been made to improve the enzyme activity, but improvements were limited and the resulting enzymes didn’t meet the requirement for practical applications (14–16).

Research has revealed that the oxidative half-reaction is very slow when using O_2_, indicating that the dioxygen is probably not the natural electron acceptor (9, 16). In fact, NicA2 was identified as a dehydrogenase using a cytochrome *c* (CycN) as its natural acceptor (17). This enzymatic electron acceptor dependence complicates NicA2’s use as nicotine catabolic system and thus sets barriers to widespread application of NicA2 for nicotine bioremediation and conversion to high-value intermediates for drug development (18, 19). In addition, the nicotine catabolic activity of NicA2 is higher than other known nicotine oxidoreductases, including Nox, NdhAB, and VppA (20–22).

Pnao and NicA both belong to the flavoenzyme family and function in the pyrrolidine pathway of nicotine mineralization (22, 23). The *k*_cat_ of Pnao for PN using O_2_ is 0.79 s^−1^, which is 127 times the aerobic *k*_cat_ of NicA2 for nicotine (24). In general, proteins sharing > 35% amino acid sequence identity are grouped into families, members of which have similar structures and carry out similar functions (25, 26). A phylogenetic tree showed that Pnao is highly homologous to NicA2, with 39.05% amino acid sequence identity (Fig. 1b and Supplementary Table 1). This indicates that the structure of NicA2 can be used as a search model in resolving the crystal structure of Pnao. As in the previous study, the substrate or co-factor preferences of similar enzymes can be changed through amino acid alteration (27–29), it is therefore possible that the substrate range of Pnao could be expanded through amino acid alteration to include nicotine.

Here, we resolved the structure of Pnao using X-ray crystallography, allowing Pnao to be used in rational-design engineering. Based on the resolved structure and size-exclusion chromatography, we found Pnao was a dimer with two flavin adenine dinucleotide (FAD) molecules bound. After identification of the substrate-binding pocket by molecular docking and alanine-scanning mutagenesis, we mutated amino acids around the substrate-binding pocket of Pnao to the corresponding residues in NicA2 to generate a nicotine-catabolizing mutant. The nicotine catabolic abilities of several Pnao mutants were optimized to the same level of NicA2 in both Tris-HCl buffer and rat serum. Through liquid chromatography/mass spectrum (LC/MS) and nuclear magnetic resonance (NMR), we identified one of the reaction products as nicotine-1’-*N*-oxide. This study details a new method for obtaining enzymes with higher nicotine-catabolizing activity using O_2_ as acceptor; furthermore, it paves a new way for developing more efficient nicotine cessation enzyme.

## Results

### High structural similarity between Pnao and NicA2 guarantees rational design

*P. putida* S16 encodes the flavin-dependent oxidases NicA2 and Pnao, which catabolize nicotine and PN, respectively, in the pyrrolidine pathway. Pnao was found to share 39.05% amino acid sequence identity with NicA2 (Supplementary Table 1), allowing us to solve the Pnao crystal structure by molecular replacement. An N-terminal truncation (30 residues) was performed to improve the quality and resolution of the protein crystals as previously reported (14). The crystal structure of PnaoΔN30 (residues 31 to 497) in complex with FAD was determined at 2.2 Å resolution using NicA2 (PDB code: 7C49) as the search model for molecular replacement (PDB code: 7E7H, Supplementary Table 4).

There were two molecules per asymmetric unit in the Pnao structure (Supplementary Fig. 1a), consistent with the molecular weight measured through size-exclusion chromatography (Supplementary Fig. 2). The Pnao dimer interface contains some hydrophobic interactions and hydrogen bonds which are between Glu134 (A) and Arg181 (B), Val132 (A) and Arg181 (B), Asn245 (A) and Asn245 (B), Arg181 (A) and Glu134 (B), His127 (A) and Glu76 (B), Glu76 (A) and Arg79 (B), and Glu76 (A) and His127 (B) (two monomers A and B, Supplementary Fig. 1b).

Two individual domains were associated with FAD and substrate binding: these were composed of 16 α-helixes, 20 β-strands, and 36 coils (Supplementary Fig. 1d). The FAD-binding domain contained a Rossmann fold with the conserved GXGXXG motif, which contacted the adenine diphosphate (ADP) portion of FAD (30) (Supplementary Fig. 1c and f). The “hotdog-like” fold and the helix bundle formed the substrate-binding domain, which contained the substrate binding site around the isoalloxazine ring of FAD (31, 32) (Supplementary Fig. 1c). Trp434 and Phe470 flank the *re*-side of the flavin’s isoalloxazine ring and constitute an “aromatic sandwich” structure, which is conserved in the flavin-dependent amine oxidase and determines substrate specificity (32) (Supplementary Fig. 1e).

The Pnao main chain could be overlaid with NicA2 with only minor deviations, suggesting that Pnao could be rationally designed through amino acid alterations. The root mean square deviation (RMSD) of this comparison was 1.143 Å; the differences were mainly caused by the different lengths and angles of several β-strands. Additionally, the fold α-helix of NicA2 (residues 453 to 455) was a coil in Pnao (residues 461 to 463) (Fig. 4a).

### Arg90 and Arg96 are essential for FAD binding and PN catabolism

Pnao contains a hydrophobic cavity for FAD placement, and 19 amino acid residues are involved in FAD binding through hydrogen bonds, attractive charges, positive-positive charges, and hydrophobic interactions (Fig. 2a and b). Whereas, the FAD-binding pocket of NicA2 has 10 residues forming 13 hydrogen bonds with the FAD (Fig. 2a), and nine of these are identical to Pnao residues (Gly67, Ala69, Glu88, Arg90, Arg96, Trp113, Val285, Phe429, and Ile463 of Pnao). Arg90 and Arg96 interact with FAD through multiple interactions. Arg90 forms one hydrogen bond and one attractive charge with FAD, and Arg96 connects with FAD through four hydrogen bonds, two attractive charges, and two positive charges (Fig. 2b). Besides these nine conserved residues between NicA2 and Pnao, Ser461 in Pnao forms two hydrogen bonds with FAD and therefore may also play an important role in FAD stabilization.

**Fig. 2.**
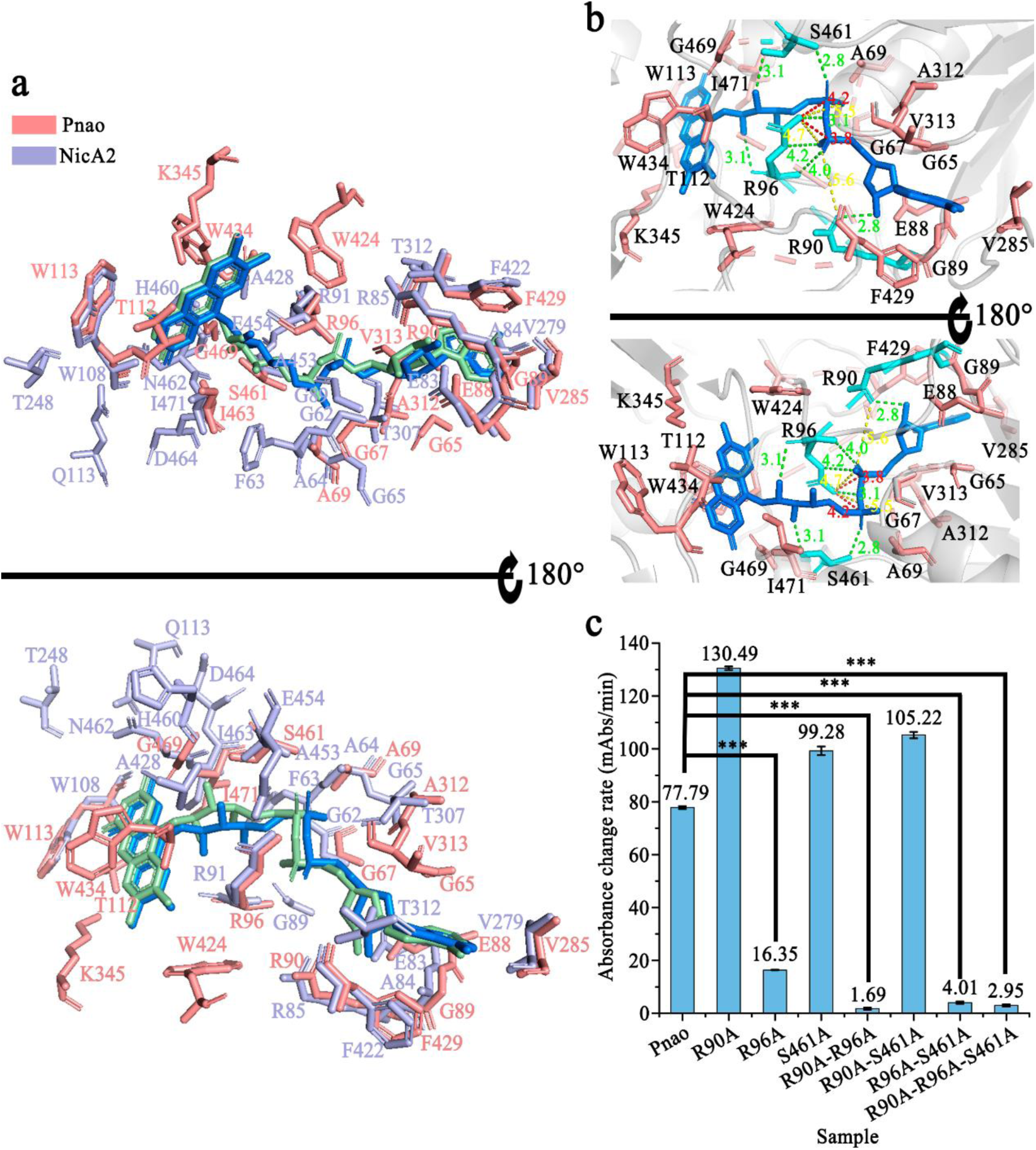
Flavin adenine dinucleotide (FAD)-binding sites in Pnao. (**a**) Comparison of amino acids in contact with FAD in Pnao (salmon) and NicA2 (light blue). **(b)** Analysis of amino acids in contact with FAD in Pnao. FAD is shown in marine, key amino acids are shown in cyan, and other amino acids are shown in salmon. The interactions between Arg90, Arg96, and Ser461 in green, yellow, and red represent hydrogen bonds, attractive charges, and positive-positive charges, respectively. **(c)** Enzyme activity of Pnao mutants relative to the wild type. Data are presented as the mean; error bars indicate standard deviation from three independent biological replicates. Three stars means p-value ≤ 0.001.

To verify the functions of the predicted residues, alanine mutagenesis was performed at positions 90 (Arg), 96 (Arg), and 461 (Ser) to generate seven mutants (Supplementary Table 2). Pnao-R96A enzymatic activity was dramatically reduced compared with other two single-point mutants (Pnao-R90A and Pnao-S461A), indicating that Arg96 is the most important residue of the three for PN catabolism (Fig. 2c). This decline in activity on PN is not due to loss of the flavin cofactor, as Pnao-R96A still purifies with FAD bound (Supplementary Table 2). Although the single mutation of R90 and S461 did not significantly reduce enzyme activity, these two residues also contributed to PN catabolism. The activity levels of Pnao-R90A-R96A, Pnao-R96A-S461A, and Pnao-R90A-R96A-S461A were extremely low (Fig. 2c). More specifically, the *k*_cat_/*K*_m_ values for PN catabolism by Pnao-R96A and Pnao-R96A-S461A dropped to 0.22% and 0.08%, respectively, of wild-type levels (Supplementary Table 3).

As shown in Supplementary Table 2, Pnao loss FAD-binding capability only if R90 and R96 were mutated at the same time (Pnao-R90A-R96A or Pnao-R90A-R96A-S461A). Mutants Pnao-S461A, Pnao-R90A-S461A, and Pnao-R96A-S461A didn’t lose FAD-binding capability, indicating that Ser461 does participate in FAD-binding, but does not play a crucial role.

Therefore, R90, R96 and S461 are all closely related to PN degrading activity, among which R96 is the most important. A single mutation on R96 can significantly reduce the enzyme activity. R90 and R96 are also essential to FAD-binding, and the mutations of both sites can cause the loss of FAD.

### Phe388 and Trp434 are critical for PN catabolism

Comparing the protein structures between NicA2-nicotine complex and Pnao, Pnao has a similar cavity at exactly the same location of the nicotine-binding pocket in NicA2. Docking between Pnao and its natural substrate, PN, was visualized using AutoDock Vina. PN could be docked into the cavity, with the active nitrogen atom facing towards the *re*-side of the FAD isoalloxazine ring (Fig. 3a). In addition, nicotine and PN molecules were located very closely in docking simulation, especially their active nitrogen atoms (Fig. 3b). Based on the high overall structural similarities of NicA2 and Pnao, we inferred that the clearly visible cavity adjacent to the isoalloxazine was the Pnao substrate-binding pocket. Additionally, Trp434 and Phe470 in this cavity form the “aromatic sandwich” conserved in the flavin-dependent amine oxidase (32), which is known to be important for proper substrate positioning.

**Fig. 3.**
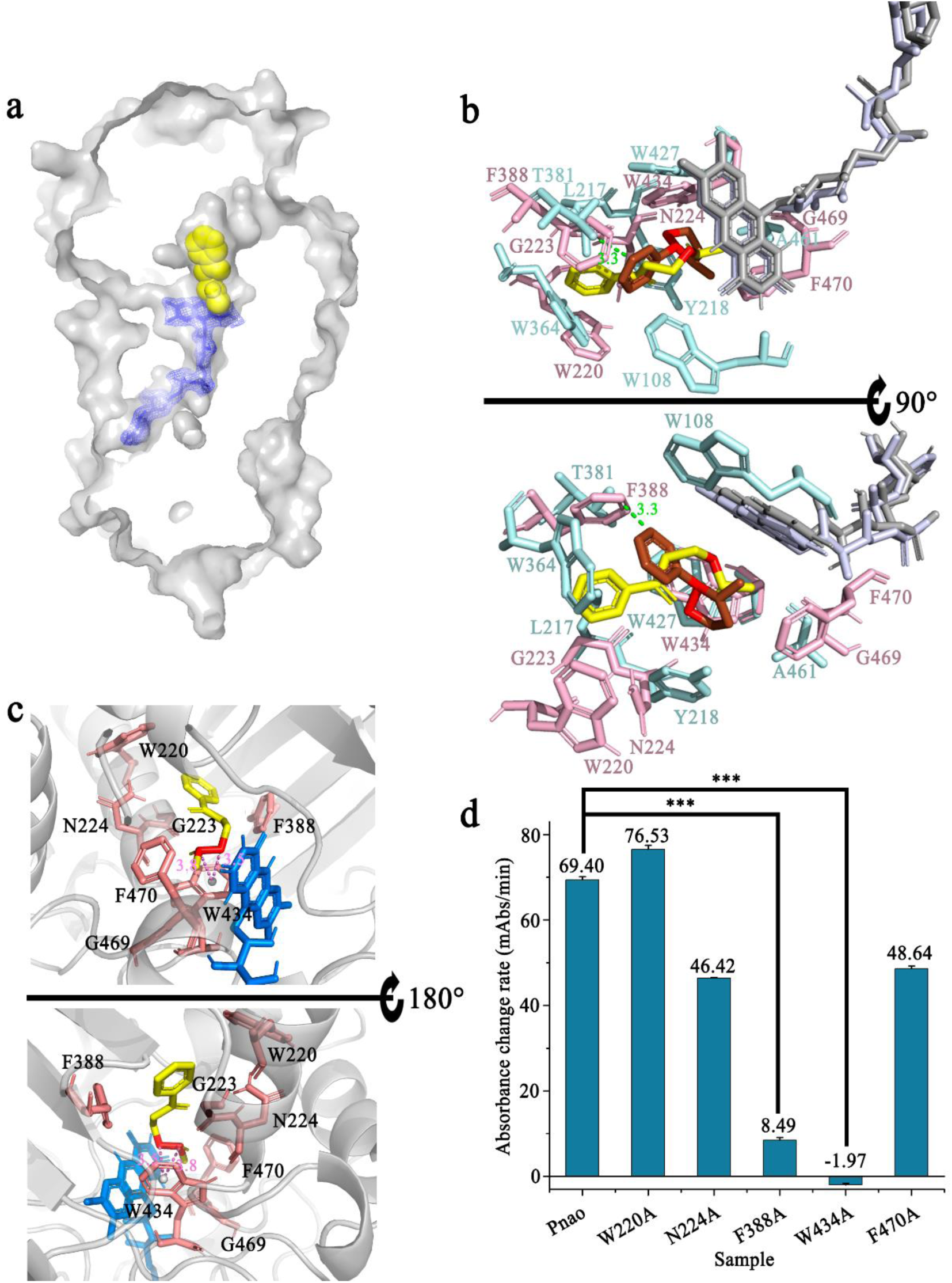
Verification of the pseudooxynicotine (PN)-binding pocket in Pnao. (**a**) AutoDock Vina showed that PN can be docked into the putative substrate-binding pocket of Pnao. A stick structure representation of flavin adenine dinucleotide (FAD) is shown in blue in a *2F_O_-F_C_* electron map, and a space-filling model of PN is shown in yellow. **(b)** Comparison of the putative PN binding site in Pnao and the nicotine binding site in NicA2. The distance between the isoalloxazine rings of Pnao and NicA2 was 0.5 Å. The distance between the active nitrogen atoms of PN (yellow) and nicotine (red) was within 3 Å. FAD in Pnao and NicA2 are shown in gray and light blue, respectively. PN is shown in green and interacting amino acids are light pink. Nicotine is shown in red and interacting amino acids are pale cyan. **(c)** Amino acids that may be in contact with PN. The active atoms (red) were adjacent to Trp434 (< 3.8 Å), an important catalytic site that is conserved between Pnao and NicA2. FAD is shown in marine, PN is shown in yellow, and relevant amino acids are shown in salmon. **(d)** Detection of PN catabolic activities of WT Pnao, W220A, N224A, F388A, W434A, and F470A. Three stars means p-value ≤ 0.001.

**Fig. 4.**
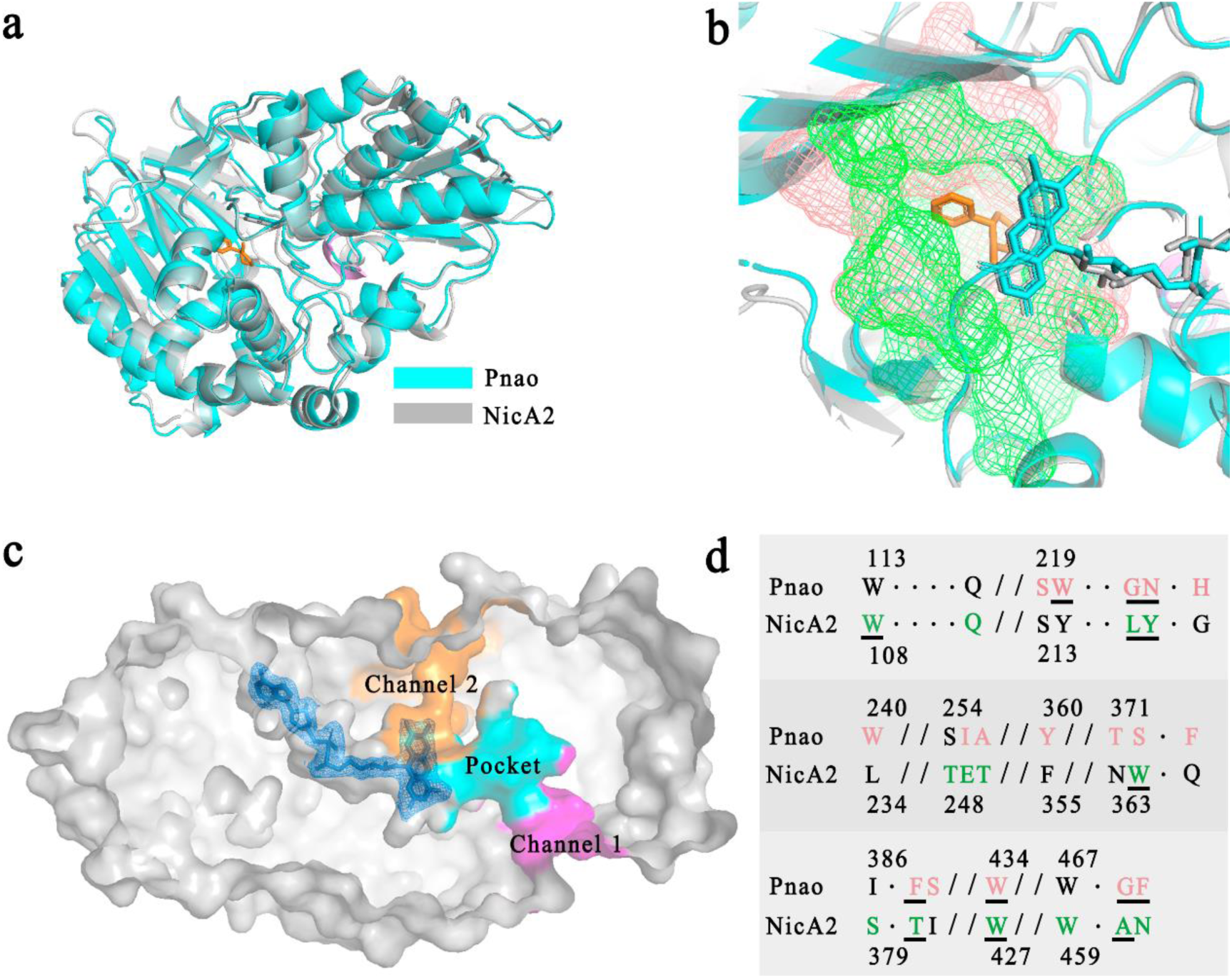
Amino acid and structure alignment of Pnao and NicA2. (**a**) Superimposed crystal structure cartoons of Pnao and NicA2 showed that they share extensive similarity, with a root mean square deviation (RMSD) value of 1.143. Pnao is shown in cyan, NicA2 is shown in gray, nicotine is shown in orange, and the α-helix of NicA2 (residues 453 to 455) is shown in violet. **(b)** The binding pockets of Pnao (salmon) and NicA2 (green) were shown overlaid with one another. **(c)** Flavin adenine dinucleotide (FAD) and putative substance channels in Pnao. The FAD molecule is colored in blue. The regions in magenta and orange are substrate channels through which PN may travel. The channel shown in magenta was considered more likely to be used by PN. **(d)** Amino acid sequence alignment of the Pnao and NicA2 binding pockets in **(b)** that are shown in detail. Amino acids in the pocket region are shown in salmon (Pnao) or green (NicA2), and underlined residues are involved in substrate binding.

This cavity is connected to the surface via two channels (Fig. 4c). Channel 2, is much narrower than channel 1, making it appear more difficult for the substrate to enter from this direction. Calculations using the program CAVER indicated that channel 2 was of lower priority than channel 1 (Supplementary Table 5). Channel 1 was therefore considered as possibly the substrate entrance tunnel, whereas channel 2 was considered as the product exit or oxygen entrance tunnel.

The amino acids involved in Pnao substrate binding were next identified from the docking model. The key residues were Trp220, Gly223, Asn224, Phe388, Trp434, Gly469, and Phe470. Notably, there is a π-σ bond between the active N-C covalent bond and Trp434 (Fig. 3c). Alanine-scanning mutagenesis was applied to five of the key amino acids (excluding Gly223 and Gly469). The mutants Pnao-F388A and Pnao-W434A almost lost PN-catabolizing activity (Fig. 3d), showing 0.23% and 0% of the wild-type activity (*k*_cat_/*K*_m_), respectively (Supplementary Table 3). This indicates that Phe388 and Trp434 are essential for PN catabolism.

### Amino acid alterations between Pnao and NicA2

As mentioned above, amino acid alteration in the active pocket based on the structure of a similar enzyme has previously been effective in altering co-factor selectivity (27, 33). Thus, here we attempted to engineer Pnao to catabolize nicotine by altering active site residues, based on the similar structures of Pnao and NicA2 (Fig. 4a and b). NicA2 contains seven amino acids involved in nicotine binding: Trp108, Leu217, Tyr218, Trp364, Thr381, Trp427, and Ala461 (Fig. 3b). Phe470, the amino acid residue consists of the “aromatic sandwich” was replaced with Asn (Fig. 4d), resulting in the loss of nicotine degrading capacity. Therefore, the adjacent residue, Gly469 was replaced with Ala. Unfortunately, all mutants resulting from alterations in the “aromatic sandwich” region, namely Pnao-G469A, Pnao-F470N, and Pnao-G469A-F470N, showed no enhancement in activity towards nicotine (Table 1).

**Table 1.** The ability of Pnao with its mutants to catabolize nicotine and pseudooxynicotine.

| **Protein** | **Nicotine**<br>**degradation (%)** | **Pseudooxynicotine**<br>**degradation (%)** |
| --- | --- | --- |
| Pnao | 6.48 ± 1.20 | 84.84 ± 0.32 |
| Pnao-G469A | 3.80 ± 1.65 | 84.01 ± 0.75 |
| Pnao-F470N | 7.92 ± 1.13 | 83.32 ± 0.47 |
| Pnao-G469A-F470N | 4.82 ± 0.96 | 52.63 ± 4.25 |
| Pnao-W220Y | 5.71 ± 1.08 | 84.31 ± 1.05 |
| Pnao-W220Y-G223L-N224Y | 11.11 ± 0.27 | 60.73 ± 3.15 |
| Pnao-W220Y-G223L-N224Y-F388T | 5.68 ± 3.20 | 8.49 ± 2.92 |
| Pnao-W220Y-G223L-N224Y-F388T-G469A-F470N | 4.00 ± 1.85 | 2.06 ± 0.21 |

Thus, residues Trp220, Gly223, Asn224, Phe388, Gly469, and Phe470, except Trp434 that is equivalent to Trp427 in NicA2, in Pnao were changed to the corresponding residues in NicA2 (Tyr214, Leu217, Tyr218, Thr381, Ala461, and Asn462) in a cumulative fashion, generating four mutants: Pnao-W220Y, Pnao-W220Y-G223L-N224Y, Pnao-W220Y-G223L-N224Y-F388T, and Pnao-W220Y-G223L-N224Y-F388T-G469A-F470N. Among the obtained mutants, Pnao-W220Y-G223L-N224Y was able to degrade nicotine (Table 1), and in an activity assay, it degraded 22.28% of nicotine in Tris-HCl buffer over 31 h, which was one-third of the capacity of NicA2 (Fig. 5a).

**Fig. 5.**
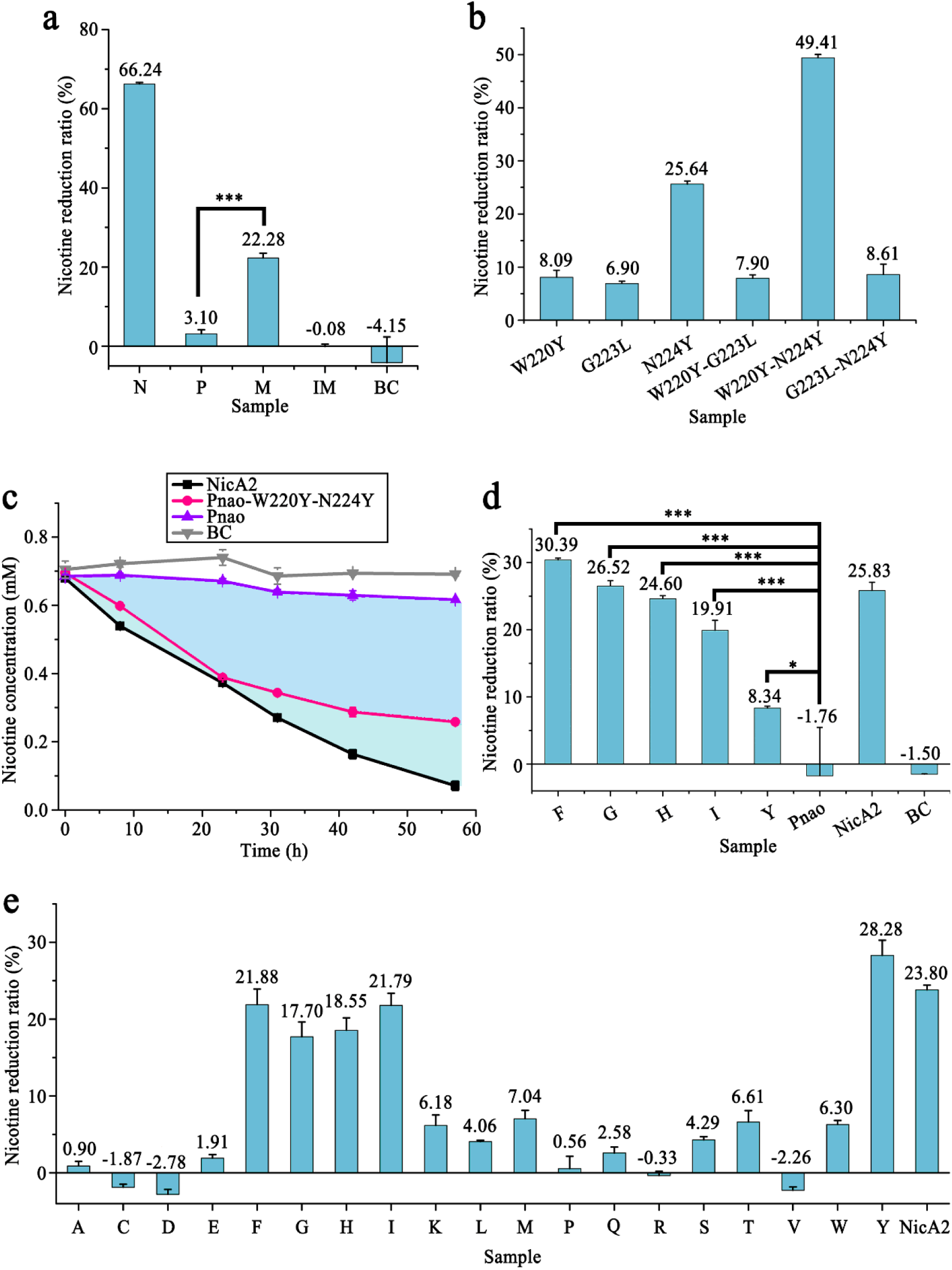
Pnao enzyme engineering. (**a**) Enzymatic activity of Pnao (P), inactive Pnao-W220Y-G223L-N224Y (IM), Pnao-W220Y-G223L-N224Y (M), and NicA2 (N) on nicotine in tris-buffer (pH 8); the blank control (BC) did not include protein. **(b)** Of the combinations of W220Y, G223L, and N224Y mutants, Pnao-W220Y-N224Y had the best nicotine catabolism performance in tris-buffer (pH 8), followed by Pnao-N224Y. **(c)** NicA2 and Pnao-W220Y-N224Y decreased concentrations of nicotine in tris-buffer (pH 8) by 89.51% and 62.76%, respectively, over three days; the enzymatic activity of Pnao-W220Y-N224Y decreased somewhat after 24 h. The BC did not include protein. **(d)** In contrast to Pnao-W220Y-N224Y (Y), nicotine concentrations were decreased by at least 20% in rat serum by Pnao-W220Y-N224F (F), Pnao-W220Y-N224G (G), Pnao-W220Y-N224H (H), and Pnao-W220Y-N224I (I). NicA2 decreased the nicotine concentration by 25.83%. The BC did not include protein. **(e)** Pnao-W220Y-N224F/G/H/I/Y decreased nicotine concentrations in Tris-buffer (pH 8) by up to 20%, which was similar to the performance of NicA2. A to Y indicate mutants generated by saturation mutagenesis of Tyr224 in Pnao-W220Y-N224Y. Dots in **(c)** and bars in **(a)**, **(b)**, **(d)**, and **(e)** represent mean values. Error bars indicate standard deviation from three independent biological replicates. Three stars means p-value ≤ 0.001 and one star means p-value ≤ 0.05.

To determine the significance of each residue in nicotine catabolism, single-point and double-point mutations were generated at Trp220, Gly223, and Asn224. Of the single-point mutants, only Pnao-N224Y can degrade nicotine (25.64% in 10 h, Fig. 5b), indicating that Tyr224 was the most crucial residue for nicotine catabolism. Addition of Tyr in place of Trp220 improved activity further, as the double-point mutant Pnao-W220Y-N224Y showed a higher nicotine degradation ratio (49.41% in 10 h, Fig. 5b), suggesting that there may be epistasis between the W220Y and N224Y mutations towards using nicotine as a substrate. The activity levels of Pnao-G223L-N224Y and Pnao-W220Y-G223L-N224Y were substantially lower than those of Pnao-N224Y and Pnao-W220Y-N224Y, indicating that Gly223 needs to be retained for nicotine catabolism (Fig. 5a and b). In a long-term time course experiment, the nicotine catabolic ability of Pnao-W220Y-N224Y was nearly the same as that of NicA2; they degraded 44.00% and 45.18% of nicotine, respectively, in 24 h. At 57 h, it increased to 62.76% and 89.51% for Pnao-W220Y-N224Y and NicA2, respectively (Fig. 5c).

### Saturation mutagenesis for better nicotine-catabolizing capability

Saturation mutagenesis was performed on Tyr224, the most crucial residue for nicotine catabolism, to enhance the nicotine-catabolizing activity of the Pnao mutant. Of the 18 resulting mutants, four exhibited nicotine catabolic activity. Specifically, Pnao-W220Y-N224F, Pnao-W220Y-N224G, Pnao-W220Y-N224H, and Pnao-W220Y-N224I degraded 21.88%, 17.70%, 18.55%, and 21.79% of nicotine, respectively, over 10 h, compared with 28.28% degradation by Pnao-W220Y-N224Y and 23.80% by NicA2 (Fig. 5e).

Pnao-W220Y-N224F was one of the best performing mutants in Fig. 5, and we further analyzed its improved nicotine degrading activity compared to wild type using anaerobic stopped-flow experiments. During substrate oxidation, the enzyme bound FAD is transformed from the oxidized to the hydroquinone (two-electron reduced) state, with a characteristic decrease in absorbance at 450 nm that corresponds with flavin reduction (34, 35). We anaerobically mixed wild-type Pnao or mutant Pnao-W220Y-N224F with a large excess of nicotine and monitored reduction of FAD over time in the stopped-flow instrument. Both enzymes reacted slowly with nicotine, taking several hundred seconds for the reactions to complete. However, from comparing the final absorbance spectra for each reaction, Pnao-W220Y-N224F was more completely reduced by nicotine than wild-type Pnao (Supplementary Fig. 5a and b). For example, mixing 10 mM nicotine with wild-type Pnao resulted in only a minor decrease in the absorbance at 450 nm whereas mixing this nicotine concentration with Pnao-W220Y-N224F reduced the absorbance at 450 nm by more than 50%. Comparing the kinetics of the reaction at lower nicotine concentrations (10 and 20 mM) was complicated by the fact that the traces for Pnao-W220Y-N224F could not be fit to an exponential function. However, traces at 50 mM nicotine, where the flavin for both enzymes were nearly completely reduced, could be fit to Equation (1) and therefore directly compared. The observed rate constant (*k*_obs_) for the reaction with Pnao-W220Y-N224F was 0.017 s^−1^ (Supplementary Fig. 5c), which is 2.7 times larger than the *k*_cat_ of NicA2 (14) and 2.4 times larger than the *k*_obs_ value of 0.007 s^−1^ observed with wild type (Supplementary Fig. 5d), confirming that the Pnao mutant reacted more rapidly with nicotine than NicA2 and wild-type Pnao.

The nicotine catabolic activity of each Pnao mutant was also tested in rat serum to explore their potential applications in nicotine cessation. The enzymatic activities of Pnao-W220Y-N224F, Pnao-W220Y-N224G, Pnao-W220Y-N224H, and Pnao-W220Y-N224I were nearly as high as NicA2 activity in rat serum over 10 h (Fig. 5d). Pnao-W220Y-N224Y did not perform as well as the other mutants.

### Nicotine-1’-*N*-oxide is one of the reaction products

We next investigated the PN catabolic activity of those nicotine catabolizing mutants. The product of PN catabolism by these mutants was SAP, as it was for the wild type (Supplementary Fig. 6). The *K*_m_ for PN of Pnao-W220Y-N224F was 81 times higher than that of wild type without a major change in the *k*_cat_ value (Supplementary Table 3). This increase in *K*_m_ results in a correspondingly lower *k*_cat_/*K*_m_ of Pnao-W220Y-N224F towards PN than wild type, indicating that there is a loss in specificity towards PN with the gain in activity towards nicotine with this mutant. The other mutants also showed increased *K*_m_ values towards PN, but the increase was comparatively mild (Supplementary Table 3). According to the *k_cat_*values, we found that the turnover frequency of Pnao-W220Y-N224I was twice that of the wild type, whereas the Pnao-W220Y-N224Y and Pnao-W220Y-N224H *k_cat_*values were ∼1.6% and 5%, respectively, of the wild-type levels (Supplementary Table 3). Overall, Pnao-W220Y-N224G and Pnao-W220Y-N224I exhibited good performance in catabolizing PN based on *k*_cat_/*K*_m_ values.

Since these Pnao mutants have activity towards both nicotine and PN, we hypothesized that they would catalyze the first two steps of the pyrrolidine pathway and catabolize nicotine to SAP. However, the liquid chromatography quadrupole time of flight mass spectrum (LC/QToF MS) analysis of the nicotine reaction products produced a mixture of compounds with *m*/*z* values of 161.1031, 163.1179 and 179.1121 solution (Supplementary Fig. 3a), none of which match that expected for SAP. The compounds whose *m*/*z* value were 161.1031 or 179.1121 match that expected for NMM and PN, respectively; however, they have different retention times and ion fragments than that of commercially-obtained NMM and PN, suggesting that the 161.1031 and 179.1121 *m*/*z* compounds may be isomers of NMM and PN (Supplementary Fig. 3a and d). We therefore isolated one of the reaction products to identify the structure of the compound. The nicotine substrate was first removed via butyl acetate extraction, then the aqueous phase was purified with high performance liquid chromatography (HPLC) twice to isolate the compound whose *m*/*z* value was 179.1121. After purification, the majority of the solution contained the product with *m*/*z* 179.1230 (Fig. 6a and b).

**Fig. 6.**
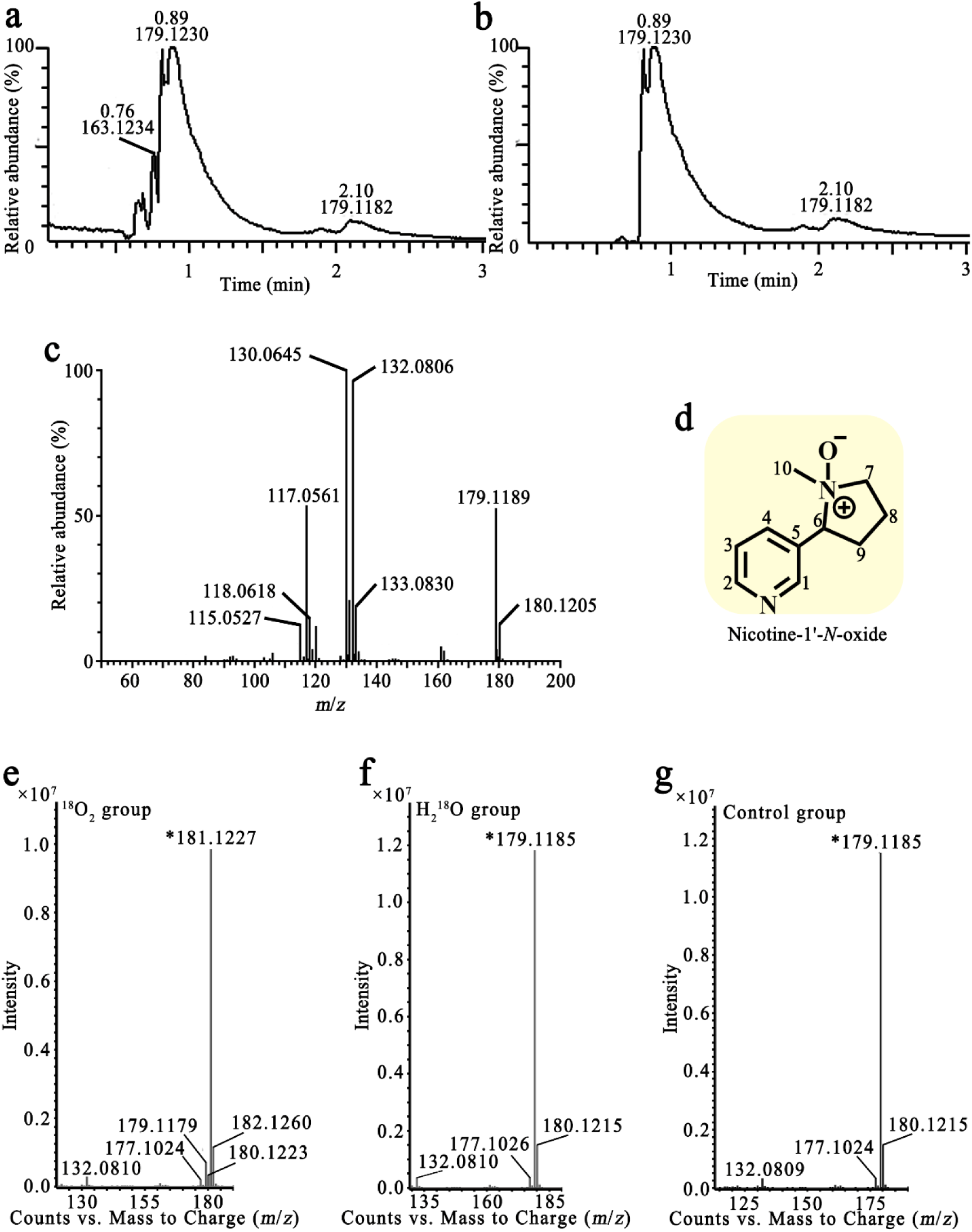
Liquid chromatography mass spectrum of the product of nicotine catabolized by Pnao mutants. (**a,b**) Total ion chromatograph **(a)** and extract ion chromatograph **(b)** of the purified product showed that the *m*/*z* value of the major product was 179.1230. **(c)** Tandem mass spectrometry spectrum of the product at *m*/*z* 179.1179. **(d)** Structural formula of the product determined by nuclear magnetic resonance analysis. **(e**–**g)** Mass spectrometry spectrum of nicotine catabolizing product under the ^18^O_2_ **(e)**, H_2_^18^O **(f)**, and natural isotope condition **(g)**. A peak at *m*/*z* 181.1227 could only be found under the ^18^O_2_ condition, suggesting that the introduced oxygen was from O_2_.

The major ion fragments at *m*/*z* of the product detected by LC/QToF MS/MS were 130.0645, 132.0806, and 179.1189 (Fig. 6c), consistent with the fragments produced by nicotine-1’-*N*-oxide (36). Electrospray ionization-MS showed an [M+H]^+^ ion at *m*/*z* 179 and the loss of 47 Da, which yielded an ion at *m*/*z* 132, corresponded to the loss of the CH_3_NH_2_ *N*-oxide from the opening of the pyrrolidine ring; a molecular hydrogen was also lost to yield a product ion at *m*/*z* 130, forming another double bond in the non-heterocyclic ring.

NMR analysis was applied to further identify the structure formula of the resulting compound. Based on NMR signals, the product was identified as nicotine-1’-*N*-oxide (Fig. 6d); the proton chemical shifts were 1.909 (m, 1H), 2.131 (m, 1H), 2.228 (m, 1H), 2.400 (m, 1H), 2.742 (s, 3H), 3.365 (m, 1H), 3.657 (m, 1H), 4.491 (t, *J* = 10.6, 1H), 7.379 (dd, *J* = 7.6/4.9, 1H), 8.082 (d, *J* = 7.9, 1H), 8.558 (d, *J* = 4.6, 1H), and 8.669 (s, 1H). The carbon chemical shifts were 19.90 (s), 28.47 (s), 53.75 (s), 69.87 (s), 74.94 (s), 122.46 (s), 129.28 (s), 138.69 (s), 149.71 (s), and 151.62 (s) (Table 2 and Supplementary Fig. 4).

**Table 2.** Nuclear magnetic resonance (NMR) signals attribution of product of nicotine catabolized by Pnao mutants.

| **Carbon position** | **Chemical shift of carbon** | **Type of carbon** | **Chemical shift of related protons (According to HSQC attribution) (ppm)** | **Multiplicity** | **Proton amounts** | **Coupling and fission values of hydrogen spectrum** | **Proton attribution related to HMBC** | **Proton attribution related to COSY** |
| --- | --- | --- | --- | --- | --- | --- | --- | --- |
| 1 | 151.62 | Quaternary | 8.669 | s | 1 | / | H-2,4,6 | / |
| 2 | 149.71 | Tertiary | 8.558 | d | 1 | 4.6 | H-1,3,4 | H-3 |
| 3 | 122.46 | Tertiary | 7.379 | dd | 1 | 7.6/4.9 | H-1,2 | H-2,4 |
| 4 | 138.69 | Tertiary | 8.082 | d | 1 | 7.9 | H-1,2,6 | H-3 |
| 5 | 129.28 | Quaternary | / | / | / | / | H-1,3,6 | / |
| 6 | 74.94 | Tertiary | 4.491 | t | 1 | 10.6 | H-1,4,7,9,10 | H-9 |
| 7 | 69.87 | Secondary | 3.657, 3.365 | m | 2 | / | H-10 | H-8 |
| 8 | 19.90 | Secondary | 2.228, 1.909 | m | 2 | / | H-7,9 | H-7,9 |
| 9 | 28.47 | Secondary | 2.400, 2.131 | m | 2 | / | H-6,7 | H-6,8 |
| 10 | 53.75 | Primary | 2.742 | s | 3 | / | / | / |

## Discussion

Most nicotine oxidoreductases are dependent on enzymatic electron acceptors such as cytochrome or pseudoazurin and exhibit poor enzyme activity when O_2_ is used as the acceptor (17, 21). This limits the breadth of applications in which nicotine catabolism can be used. In addition, mutations in NicA2, the most effective and widely studied nicotine catabolizing enzyme, have shown very limited improvement in reaction activity using O_2_ (14–16). To obtain a suitable enzyme for broader applications of nicotine catabolism, a new research direction was proposed and executed here. Pnao, which has higher efficiency than NicA2 when using O_2_ as the electron acceptor, was engineered to catabolize nicotine, and one of the reaction products was identified as nicotine-1’-*N*-oxide.

The origin of the introduced oxygen to nicotine was analyzed via isotopic oxygen labeled O_2_ gas and water. The ion fragments at *m*/*z* of product produced in the presence of ^18^O_2_ detected by LC/QToF MS were 132.0810, 177.1024, 179.1179, 180.1223, 181.1227, and 182.1260 (Fig. 6e). The *m*/*z* 181.1227 and 182.1260 signals are two Daltons larger than the 179.1185 and 180.1215 signals observed in the control reaction, indicating that an oxygen atom from O_2_ was incorporated in the product. In contrast, the spectral peaks observed when using H_2_^18^O (Fig. 6f) matched that of the control sample (Fig. 6g). Together, these results indicate that the introduced oxygen atom of nicotine-1’-*N*-oxide came from O_2_ rather than H_2_O.

The oxidation of substrate by Pnao in the reductive half reaction does not use O_2_ directly; O_2_ oxidizes the reduced flavin in the oxidative half reaction to return the reduced flavin to the oxidized state, producing H_2_O_2_ as a byproduct. It is known that tertiary amines like nicotine can react with H_2_O_2_ to form amine-*N*-oxides (37), and LC/MS confirmed that nicotine could be converted to nicotine-1’-*N*-oxide in solution upon adding H_2_O_2_ (Fig. 7a). Nicotine-1’-*N*-oxide could not be detected in the nicotine reaction products produced by Pnao mutant when catalase was added (Fig. 7b and c), suggesting that nicotine-1’-*N*-oxide was a byproduct of the H_2_O_2_ produced by Pnao mutant. However, electrons must be removed from something–most likely nicotine–in order to generate H_2_O_2_ during catalysis, which would require production of another product. Apart from nicotine and nicotine-1’-*N*-oxide, other generated chemicals such as the one whose *m*/*z* value was 161.1031 (the same value as *N*-methylmyosmine), were detected (Supplementary Fig. 3a). And it was also detected in the nicotine catabolic products of Pnao mutant when catalase added (Supplementary Fig. 3b). Furthermore, the amount of nicotine-1’-*N*-oxide produced corresponds to only 17% of the amount of nicotine consumed in the reaction, indicating that there indeed must be other products produced in the reaction that we were unable to identify (Fig. 7d), leaving an open question about the nicotine catalytic mechanism used by these mutants (Fig. 7e).

**Fig. 7.**
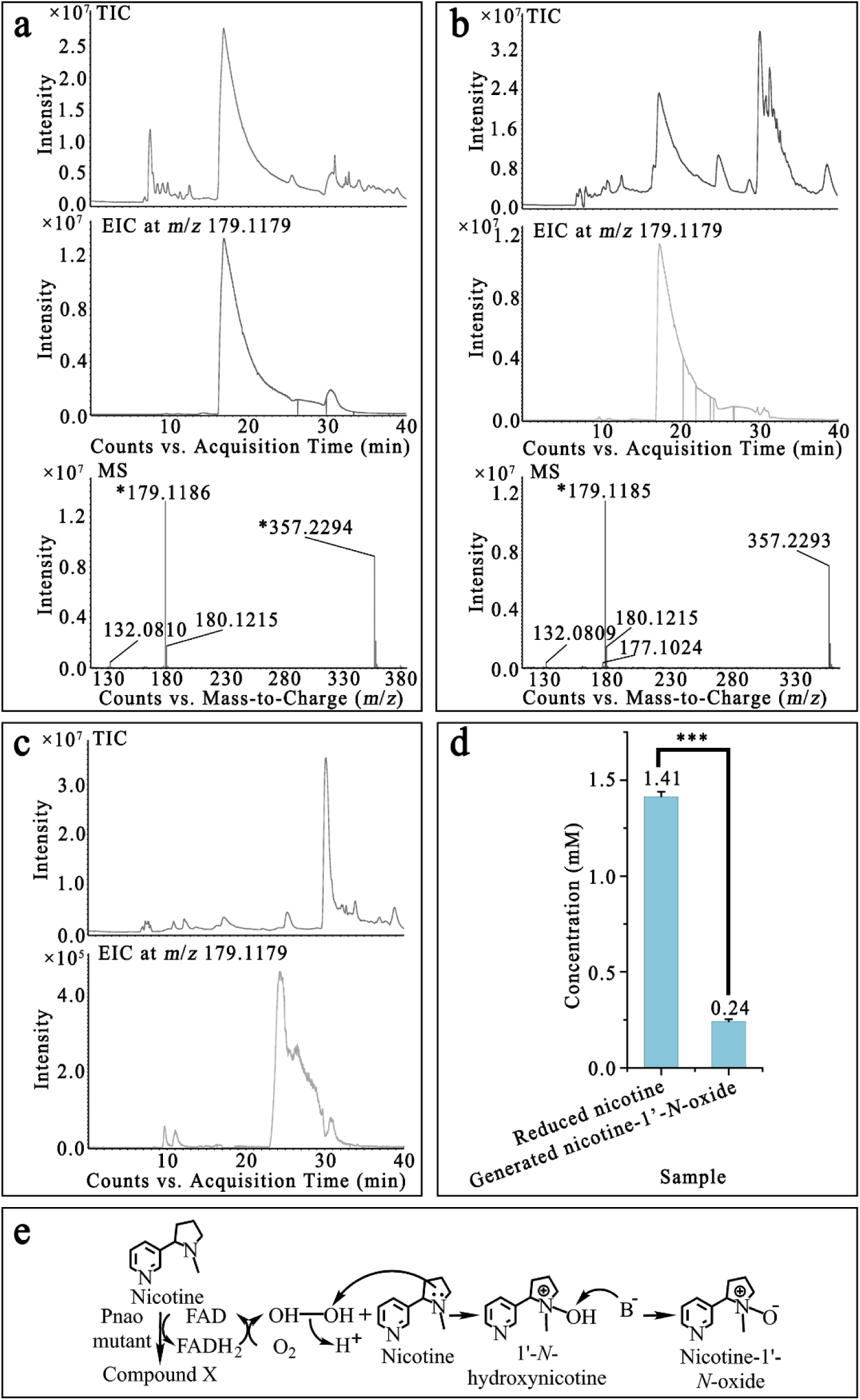
Proposed nicotine catabolic mechanism of Pnao mutant. (**a,b**) Liquid chromatography quadrupole time of flight mass spectrum (LC/QToF MS) of nicotine catabolized by H_2_O_2_ **(a)** and Pnao mutant **(b)** indicated that nicotine could be oxidized by H_2_O_2_. **(c)** Nicotine-1’-*N*-oxide was not present in the reaction solution when nicotine was incubated with Pnao mutant and catalase. **(d)** Concentrations of reduced nicotine and generated nicotine-1’-*N*-oxide in the reaction solution of nicotine and Pnao mutant. Error bars indicate standard deviation from three independent biological replicates. **(e)** Proposed nicotine catabolic mechanism of Pnao mutant. TIC refers to total ion chromatograph, EIC refers to extract ion chromatograph, and MS refers to mass spectrum. Three stars means p-value ≤ 0.001.

To further increase nicotine catabolic activity, mutational screens could be conducted that focus on the active sites (38), the FAD-binding pocket (39), the oxygen-binding area (40, 41), and the channels for substrate entrance and product exit (14, 42). NicA2 for example, nine bulky residues around the product exit channel that reduce the release speed of PN were previously mutated to alanine or valine based on the α-helix or β-strand folding preference, and the *k_cat_* value of the mutant increased to 3.7 times that of the wild-type protein (14). Therefore, the nicotine catabolic activity of the mutant might be further enhanced by mutating 11 residues of channel 2 in Pnao (the putative product exit): His226, Lys345, Pro392, Ile395, Ile397, Asn398, Tyr420, Tyr422, Trp424, Asn425, and Leu426.

Here, we summarized a strategy for obtaining high activity of O_2_-depending nicotine-catabolizing enzyme. Two empirical rules should be followed in such efforts based on the findings of Pnao engineering in the present study. First, a low RMSD between the structures of the selected enzyme and NicA2 is required as a foundation; protein structure determines function, and proteins with similar structures often have analogous functions (43). Superoxide dismutases (SODs) have a range of metal specificities. The Mn SOD and Fe SOD that binding manganese and iron ions, respectively, in *Staphylococcus aureus* exhibit extensive structure similarities, and enzyme engineering can alter cofactor specificity (27). Second, amino acid alteration around the substrate-binding pocket is a useful and feasible strategy. Mutating the substrate binding region to enlarge the binding pocket is efficient to extend substrate range, which has been shown in *Aspergillus niger* monoamine oxidase N (MAO-N). MAO-N was engineered to catalyze benzhydrylamine, which is two times larger than its natural substrate (benzylamine), after several random mutations around the substrate-binding pocket that enlarged the active site (29). In Pnao, steric hindrance may determine the substrate specificity. When Trp220, which has a very large indole group, was mutated to a smaller residue (tyrosine) and Asn224 was mutated to glycine in a computer simulation (Supplementary Fig. 7c), and the sidechains of phenylalanine, histidine, isoleucine, and tyrosine rotated to a suitable angle, the substrate-binding pocket was enlarged, which allows the nitrogen atom of nicotine pyrrolidine to turn to a more logical direction, facing the *re*-side of the FAD isoalloxazine ring (Supplementary Fig. 7b and d–f).

Such a strategy could be applied to other similar enzymes like L-amino-acid oxidase (LAAO), polyamine oxidase (PAO), and MAO. MAO plays important roles in the oxidation of biogenic and xenobiotic amines, such as neuroactive serotonin, norepinephrine, and dopamine (44, 45). MAOA and MAOB are enzymes from human and similar in structure and amino acid sequence to NicA2. They could be used in nicotine cessation with the tremendous advantage in minimizing immunological rejection if they could be engineered to adequately catabolize nicotine like Pnao (Supplementary Fig. 8 and Supplementary Fig. 9).

In summary, we reported the structure of Pnao and verified the FAD and substrate-binding pockets via alanine-scanning mutagenesis. Based on structural similarity with NicA2, amino acid alterations around the substrate-binding pocket allowed Pnao to catabolize nicotine. Steric hindrance was proposed as an explanation for Pnao substrate specificity. In the future, catabolic activity of Pnao-W220Y-N224Y could be enhanced by mutating the amino acids around the FAD-binding pocket. These strategies could also be applied to more enzymes, allowing wider applications on nicotine catabolism and forging a new direction in nicotine cessation research.

## Materials and Methods

### Chemical reagents

L-(−)-Nicotine and 4-(methylamino)-1-(3-pyridyl)-1-butanone dihydrochloride (pseudooxynicotine) were purchased from Sigma-Aldrich (USA) and Toronto Research Chemicals (Canada). Rat serum was obtained from Sbjbio Life Sciences (China). Crystallization reagents were purchased from Hampton Research (USA). All other reagents were obtained commercially.

### Protein purification

Gene *pnao* was cloned into the plasmid pET-28a (Novagen) between the *Nco*I and *Xho*I sites. Site mutations and protein truncations were conducted via whole-plasmid PCR amplification and *Dpn*I digestion (46). Recombinant plasmids were validated with Sanger sequencing (Personalbio Technology, China), then transformed into *Escherichia coli* strain BL21(DE3). Cells containing the plasmids were cultivated in Luria broth (LB) medium at 37°C with 200 rpm shaking until the OD_600_ reached 0.8. After cultures were cooled to 16°C, protein expression was induced by incubation with 0.3 mM isopropyl β-d-1-thiogalactopyranoside (IPTG) overnight. Cells were harvested by centrifugation at 4,000 × *g* for 20 min at 4°C; the precipitate was resuspended in balance buffer (25 mM Tris-HCl [pH 8.0], 300 mM NaCl, 20 mM imidazole, 1 mM phenylmethylsulfonyl fluoride [PMSF], and 10 mM β-mercaptoethanol) prior to nickel nitriloacetic acid (Ni^2+^-NTA) affinity chromatography (BBI Life Sciences, China). Cells were lysed twice at 800 bar with a high-pressure homogenizer (ATS Engineering, Canada) at 4°C, followed by centrifugation at 10,000 × *g* for 1 h at 4°C. The supernatant was loaded onto the Ni^2+^-NTA column after filtration (with 0.22 μm pore size) (BBI Life Sciences, China) to obtain the target protein through gradient elution. For enzyme assays, enzyme assay buffer (25 mM Tris-HCl [pH 8.0], 200 mM NaCl, and 5 mM dithiothreitol [DTT]) was applied as the elution buffer and ultrafiltration was performed to concentrate the enzyme. For crystallization, the eluate was loaded onto size-exclusion chromatography columns (Superdex 200 Increase 10/300 GL, Cytiva Life Sciences, USA) to separate polymers and replace the buffer in which proteins were suspended. The contents of all fractions were confirmed with sodium dodecyl sulfate-polyacrylamide gel electrophoresis (SDS-PAGE). Isolated proteins were frozen in liquid nitrogen and stored at −80°C until use.

### Crystallization, data collection, and structure determination

Pnao crystals were grown in a 1:1 (v/v) mixture of Pnao protein solution (20 mg mL^−1^) and a reservoir solution (0.2 M lithium sulfate monohydrate, 0.1 M BIS-TRIS [pH 6.5], and 25% w/v polyethylene glycol 3350) by hanging-drop vapor diffusion at 20°C. Crystals were transferred to reservoir solution containing 25% (v/v) glycerol, then cryocooled in liquid nitrogen.

Crystal diffraction datasets were collected with the BL19U1 beamline at the Shanghai Synchrotron Radiation Facility using a Platus M6 detector at a wavelength of 0.97918 Å at 100 K. All data were processed and scaled using the *HKL*-3000 program (47). Structures were determined using the molecular replacement method with *Phaser* in CCP4 (48), using NicA2 (PDB code: 7C49) as the searching model. After model building with *Coot* (49), and refinement with *REFMAC* in CCP4 (50), small molecules were placed into the model based on the Fourier difference map and refined using the geometric restraints output by Restrain in CCP4 (51). All structural figures were drawn in PyMOL. Crystallographic statistics are shown in Supplementary Table 4.

### Enzyme assay

Two methods were used to characterize enzymatic activity: UV-2550 spectrometry (Shimadzu, Japan) and high-performance liquid chromatography (HPLC) on an Agilent 1200 instrument (Agilent, USA) (24). The reaction was performed in 25 mM Tris-HCl buffer (pH 8.0) at 30°C. The rate of A_230_ change within 1 min was monitored and analyzed via UV-2550 spectrometers to compare the enzymatic activity in solutions with 0.1 mM PN and 0.1 mg mL^−1^ enzyme. Enzyme kinetics were determined based on substrate-enzyme reactions at 0.1 mg mL^−1^ enzyme and a range of initial substrate concentrations. Final substrate concentrations were measured 2 min after the reactions started via HPLC at 260 nm after quenching with 10 μL of 37% HCl. The Eclipse XDB-C18 column was used, which was 5 μm and 4.6 × 150 mm (Agilent, USA). The mobile phases were 95% ultrapure water with 0.1% (v/v) formic acid and 5% methanol, and the mobile flow rate was 0.3 mL min^−1^.

To determine the catabolic activity of Pnao mutants (1 mg mL^−1^) in Tris-HCl buffer or rat serum, 1 mM nicotine or 0.5 mM PN was added to each solution. The final concentration of nicotine or PN after incubation for several hours at 30°C was analyzed with HPLC. Nicotine reactions were stopped by adding an equal volume of ethyl acetate, vortexing for 1 min, then centrifuging at 13,800 × *g* for 1 min to harvest the organic phase. PN reactions were quenched by adding 10 μL of 37% HCl.

### Pnao mutation

The mutagenesis was performed by pEASY-Uni Seamless Cloning and Assembly Kit (TransGen, China). The primers are listed in Supplementary Table 6. The mutants of *pnao* genes were inserted into pET28a and transferred into *E. coli* BL21(DE3). The enzyme activity was detected as mentioned above.

### Product identification

To identify the product of Pnao mutant nicotine catabolism, 0.2 mg mL^−1^ protein and 5 mM nicotine were mixed in 25 mM Tris-HCl buffer (pH 8.0) and incubated at 30°C for three days. The remaining nicotine was removed by butyl acetate extraction five successive times, then the product was concentrated and separated by HPLC on the Agilent 1200 (Agilent, USA). Fractions of main peaks were collected and identified through LC/MS in positive ion mode on the Xevo G2-XS QToF (Waters, USA). The purified products were collected and mixed for another separation by HPLC and identified through LC/MS before analysis via NMR on the AVANCE NEO at 700 M Hz (Bruker, Germany). The Eclipse XDB-C18 column was used, which is 5 μm and 4.6 × 150 mm (Agilent, USA). The mobile phases of first-step HPLC separation were 95% ultrapure water with 0.1% (v/v) formic acid and 5% methanol; the second-step HPLC separation phases were 95% ultrapure water with 0.05% (v/v) ammonium hydroxide (25%) and 5% methanol. The flow rate was 0.3 mL min^−1^.

### Isotopic oxygen analysis

^18^O labeled oxygen gas and water were the reaction medium in figuring out where the introduced oxygen of nicotine catabolizing product from. In ^18^O_2_ group, oxygen in the reaction buffer was removed by balanced in the anaerobic incubator (Punmicro Scientific, China) for several days before ^18^O_2_ inflation. In H_2_^18^O group, the Tris-HCl (pH 8.0) buffer was prepared with H_2_ ^18^O. In control group, there was no ^18^O replacement. 0.6 mg mL^−1^ enzyme with 2 mM nicotine was incubated at 30°C for 3 days before mass spectrometry on 6545 Q-TOF LC/MS system (Agilent, USA) in positive mode. The Eclipse XDB-C18 column was used, which is 5 μm and 4.6 × 150 mm (Agilent, USA). The mobile phases were 95% ultrapure water with 0.1% (v/v) formic acid and 5% methanol and the flow rate was 0.3 mL min^−1^.

### FAD-binding ratio

The FAD concentration was analyzed by high performance liquid chromatography (HPLC) described before (24). Protein concentration was detected by Nano Drop (Thermo Scientific, USA) and then held the protein in boiled water for 10 min. The denatured protein was removed after centrifuged at 13,800 × *g* for 10 min. The supernatant containing FAD was analyzed by HPLC on the Agilent 1200 (Agilent, USA). The Eclipse XDB-C18 column was used, which is 5 μm and 4.6 × 150 mm (Agilent, USA). The mobile phases were 70% 10 mM ammonium sulfate and 30% methanol. The flow rate was 0.5 mL min^−1^. The wavelength was 266 nm.

### Stopped-flow

Stopped-flow experiments for Pnao variants were carried out as described previously (52) in 40 mM HEPES-KOH pH 7.5, 100 mM NaCl, 10% glycerol at 4°C using a TgK Scientific SF-61DX2 KinetAsyst stopped-flow instrument. Pnao solutions were made anaerobic in glass tonometers by cycling with vacuum and anaerobic argon (53). The tonometer was then loaded onto the instrument and the enzyme solution was mixed with buffer containing 20, 40, or 100 mM nicotine (concentration before mixing in the instrument) that had been sparged with argon to achieve anaerobiosis, and the subsequent reaction was monitored using the instruments multi wavelength CCD detector. Reaction traces at 450 nm were plotted in KaleidaGraph, and the traces using 50 mM nicotine after mixing were fit to Equation (1). In Equation (1), ΔA is the kinetic amplitude, *k*_obs_ is the apparent first order rate constant and A_∞_ is the absorbance at the end of the reaction.

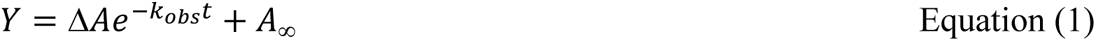

### Structure and sequence analysis

BLASTP searches and phylogenetic tree construction were carried out in MEGA-X using ClustalW and the maximum likelihood method. Ligand docking was performed in AutoDock Vina with default parameters (54), and interactions between ligands and enzymes were analyzed in Discovery Studio 2016. The simulative structure of Pnao mutants in silicon were generated via rotamer on UCSF Chimera version 1.13.1 (55). The Pnao cavity volume was calculated with CAVER 3.0.3 in PyMOL (56).

### Statistical analysis

Data analysis and graphing were conducted in OriginPro 9.

## Data availability

All data are available in the main text or the supplementary materials. The coordinates for the structure of Pnao have been released in the Protein Data Bank under PDB ID 7E7H.

## Acknowledgements

This project was funded by the National Key R&D Program of China 2021YFA0909500 (H. T.), the National Natural Science Foundation of China 32030004 (H. T.), the Oceanic Interdisciplinary Program of Shanghai Jiao Tong University SL2020MS027 (H. T.), and the National Institutes of Health R15GM139069 (F. S.)

## Contributions

H. H., Z. X., P. X., and H. T. outset and designed experiments. Z. X. and Z. Z. performed experiments. H. H., Z. X., and P. S. anaylzed data. H. T. and F. S. received projects, contributed reagents and materials. Z. X., H. H., and H. T. wrote the paper. All authors importantly discussed and revised the manuscript. All authors commented on the manuscript before submission. All authors read and approved the final manuscript.

